# Sugar-induced cell death in exponentially growing yeast depends on the functionality of the nonoxidative branch of the pentose phosphate pathway

**DOI:** 10.64898/2026.02.07.704583

**Authors:** Airat Ya. Valiakhmetov

## Abstract

Sugar-induced cell death (SICD) remains an intriguing but poorly studied phenomenon in the physiology of *Saccharomyces cerevisiae*. Recently, it was shown that SICD development largely depends on the redirection of glucose fluxes between glycolysis and the pentose phosphate pathway (PPP). In particular, inhibition of glycolysis by iodoacetamide (IAA) was observed to reduce SICD levels. This study is devoted to further investigation of the relationship between SICD and the functionality of the two PPP branches. It was shown that deletion of the *ZWF1* gene does not affect the decrease in SICD levels in IAA-treated cells. This allows us to conclude that the oxidative branch of the PPP is not involved in the suppression of SICD/ROS. Deletion of the *GLR1* gene and attenuation of the *TRR1* gene also did not restore SICD levels in cells after IAA treatment. The obtained results indicate that the level of reduced glutathione or thioredoxin does not affect SICD genesis. The addition of 5 mM ribose-5-phosphate (R5P) to the incubation medium led to suppression of SICD by 79%. At the same time, the addition of 5 mM ribose + 5 mM Pi suppressed SICD by only 20%. Suppression of SICD by 5 mM R5P in the *Δpho3* strain (83%) excludes the mechanism of extracellular dephosphorylation of R5P to ribose, its subsequent transport into the cell, and re-phosphorylation inside the cell. Furthermore, more than 70% suppression of SICD in the *Δend3* strain with 5 mM R5P excludes endocytosis as a mechanism of R5P import into the cell. The observed effect of R5P can be explained by the moonlighting function of some unknown protein. Thus, SICD development in *S. cerevisiae* cells depends on the final product of the non-oxidative PPP—R5P.

## Introduction

SICD (sugar-induced cell death) was first described in 1991 (Granot and Snyder 1991). The phenomenon consisted of the death of stationary-phase cells incubated only with glucose in the absence of other essential Vcompounds. A necessary condition for SICD development was the phosphorylation of glucose or fructose (Granot and Snyder 1993; Granot and Dai 1997). This led to ROS accumulation and subsequent apoptosis (Granot, Levine, and Dor-Hefetz 2003). SICD was also observed under strictly anaerobic conditions (Yoshimoto *et al*. 2009). Later, SICD development was shown in exponentially growing yeast. In this model, SICD in the form of primary necrosis occurred much faster (minutes vs. days), was accompanied by increased numbers of ROS-overloaded cells, and affected only cells in the S-phase of the cell cycle (Valiakhmetov *et al*. 2019). This necrosis was extremely sensitive to extracellular pH and membrane potential, and independent of mitochondrial respiratory chain functionality.

Specifically, dissipation of ΔpH or ΔΨ led to almost complete suppression of SICD (Bidiuk, Alexandrov, and Valiakhmetov 2021). Although SICD was long considered absent in pathogenic *Candida* species, recent results first showed temperature-dependent SICD development in several *Candida* species (Parbhudayal and Cheng 2025a). The same authors also published the most comprehensive review of SICD to date (Parbhudayal and Cheng 2025b), covering all aspects of this phenomenon. Recently, it was shown that SICD in exponentially growing *S. cerevisiae* critically depends on the redirection of glucose fluxes between glycolysis and the pentose phosphate pathway (PPP) (Valiakhmetov 2025). Disruption of glycolysis by deletion of the *HXK2, TDH3, TPS2*, or *PFK1* genes led to suppression of SICD. To explain these data, it was hypothesized that under SICD-inducing conditions, the PPP does not function fully, leading to NADPH deficiency and, consequently, ROS excess. The latter leads to SICD. These data, combined with the fact that SICD occurs only in S-phase cells (when PPP activity is sharply increased), raised the question of the role of PPP in SICD development.

PPP is the second most important metabolic pathway for glucose utilization after glycolysis and consists of two branches (Stincone *et al*. 2015; Bertels, Fernández Murillo, and Heinisch 2021). The irreversible oxidative branch uses glucose-6-phosphate to produce ribulose-5-phosphate while reducing two molecules of NADP^+^ to NADPH. NADPH is the main reducing cofactor for antioxidant enzymes (Pollak, Dölle, and Ziegler 2007). A high NADPH/NADP^+^ ratio is essential for coping with oxidative stress. The reversible non-oxidative branch intercepts glycolytic intermediates fructose-6-phosphate (F6P) or glyceraldehyde-3-phosphate (GAP) to produce the main PPP product—ribose-5-phosphate (R5P), necessary for nucleic acid synthesis. Thus, each PPP branch can function at different intensities to meet specific cellular needs. Redirection of glucose flux from glycolysis to the PPP under oxidative stress was demonstrated upon inactivation of triose phosphate isomerase (EC 5.3.1.1) and glyceroaldehyde-3-phosphate dehydrogenase (EC 1.2.1.12 GAPDH) in *S. cerevisiae* (Ralser *et al*. 2007). This led to an increase in the NADPH/NADP^+^ ratio, allowing cells to successfully resist oxidative stress. The rise in PPP metabolite concentrations proves that PPP was indeed activated. In contrast, *Δzwf1* and *Δgnd1* strains, unable to regenerate NADP^+^ via the oxidative PPP, showed high sensitivity to acrolein-induced oxidative stress (Kwolek-Mirek *et al*. 2025).

This study aimed to investigate the role of PPP in SICD development in *S. cerevisiae*, particularly to determine which PPP branch and by what mechanism suppresses SICD (Valiakhmetov 2025).

## Materials and methods

### Culture growth

Strains *Δzwf1, Δglr1, Δpho3, Δend3*, and their parental strain BY4742 were obtained from the Euroscarf collection. Strain TRR1 with decreased abundance by mRNA perturbation, *TRR1*_*DAmP*,_ was obtained from Horizon Discovery (Horizon Discovery, Cambridge, MA, USA). The culture grew on the standard YPD medium (Applichem, Darmstadt, Germany) for 15–17 h (mid-exponential phase) and was twice washed with distilled water and once with MilliQ water. Yeast cells were pelleted and suspended (1 g/10 mL w/w, 1.45 × 10^8^ cells/mL) in MilliQ water.

### 1,2,3-Dihydrorhodamine (DHR) Staining

To estimate the number of cells with elevated intracellular ROS levels, we used DHR (Sigma-Aldrich, St. Louis, MO, USA). 6 µL of 0.5 mg mL^-1^ DHR in DMSO was added to 300 µL cell suspension in water. The samples were incubated in a ThermoMixer (Eppendorf, Hamburg, Germany) for 1 h at 30°C (27 °C for *TRR1DAmP*). The cells were pelleted at 13,000× g for 1 min, and the supernatant was discarded. The cells were washed once with 0.5 mL of water, and the cell pellet was suspended in 300 µL of water. Another 300 µL of cell suspension in water without the addition of DHR was subjected to all the above steps to obtain cells for propidium iodide (PI) staining (see below).

### SICD Assay and Flow Cytometry

SICD assay for DHR-stained and unstained cells was done in a parallel series of tubes. Cells obtained as described above were distributed into 6 tubes. Nothing was added to the first tube (50 μL) —this was the control. All other test tubes were filled with cell suspension, test substances, and glucose in volumes that yielded the desired concentration in a final volume of 50 µl. The minimum cell suspension volume was 44 µl. The final glucose concentration was always 100 mM. The samples were incubated in a ThermoMixer (Eppendorf, Hamburg, Germany) at 30 °C for 1 h (27 °C for *TRR1*_*DAmP*_). To proceed to flow cytometry to the samples with DHR-stained cells, 1 mL of MilliQ water was added. PI staining was used to determine the percentage of dead cells. To the samples with not DHR-stained cells, 150 µL of 4 µg mL^-1^ PI (Sigma-Aldrich, St. Louis, MO, USA) was added, and after brief (1–2 min) incubation at RT, 0.8 mL of MilliQ water was added. All samples were kept on ice. A total of 100,000 cells were counted at each experimental point using a NovoCyte Flow cytometer (Agilent, Santa Clara, CA, USA). DHR-stained cells were detected using 488 nm for excitation and 530/30 nm for emission. PI-stained cells were detected using 488 nm for excitation and 572/28 nm for emission. All assays were repeated three times, and the mean results are presented.

### Iodoacetamide (IAA) assay

Cells were prepared as in the *Culture growth* and *SICD Assay* sections and incubated with 1, 1.5, and 2 mM of IAA. After 20 min incubation at 30 °C (27 °C for *TRR1*_*DAmP*_), to induce SICD, 2 M glucose solution was added to all tubes (final [C] = 100 mM). The samples were incubated at 30 °C for 1 h (27 °C for *TRR1*_*DAmP*_). Then, samples proceed to flow cytometry. Due to the instability of the aqueous solution of IAA, stock solutions were prepared 20 minutes before application.

### Glucose uptake assay

To assess glucose concentration, the GOD-PAP colorimetric method was employed using the “Glucose DDC” kit (Diakon-DC, Pushchino, Russia) according to the instructions.

## Results and Discussion

Redirection of carbon fluxes from glycolysis to the PPP upon glycolysis disruption has been repeatedly confirmed (Ralser *et al*. 2006, 2007; Grüning *et al*. 2011, 2014). Previous analysis of SICD suggested that suppression of this phenotype upon partial inhibition of glycolysis with iodacetamide (IAA) might involve a redirection of carbon flux from glycolysis into the PPP (Valiakhmetov 2025). We hypothesized that impairment of glycolytic flux with 2 mM IAA, a specific inhibitor of GAPDH (Lind *et al*. 1998; Grant 2008), could lead to accumulation of sugar phosphates upstream of the block, particularly GAP and F6P, which might then be diverted into the oxidative branch of the PPP. In this scenario, increased NADPH production would support glutathione reductase activity (EC 1.8.1.7, Glr1p), thereby enhancing glutathione-dependent ROS scavenging and ultimately suppressing SICD. The accumulated GAP can enter the PPP through the non-oxidative branch, generating ribose-5-phosphate (R5P) to support dNTP synthesis. Triose phosphates can also be metabolically recycled to F6P and G6P through the reversible non-oxidative PPP. This prompted us to examine both branches of the PPP experimentally. First, we examined whether ZWF1 is required for the IAA-mediated suppression of SICD/ROS. However, GAPDH inhibition by 2 mM IAA did not affect SICD/ROS dynamics in the *Δzwf1* strain (Fig. 1). This indicates that the oxidative branch of the PPP is not involved in the observed suppression of SICD/ROS. But the oxidative branch is not the only NADPH source in the cell. Isocitrate dehydrogenase (EC 1.1.1.42) and aldehyde dehydrogenase (EC 1.2.1.4) can also contribute to maintaining the NADPH pool (Pollak, Dölle, and Ziegler 2007).

**Fig 1.**
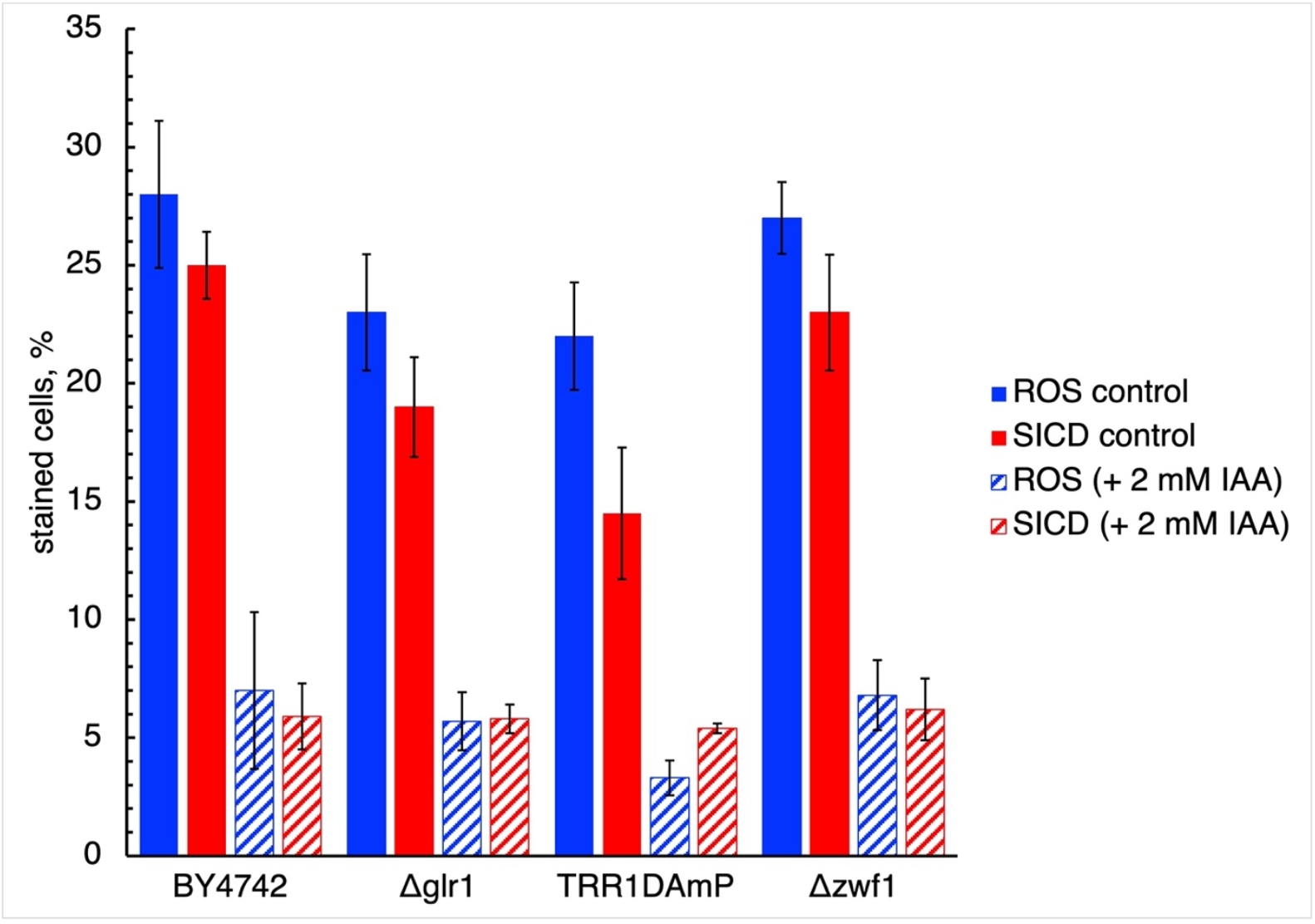
Effect of Tdh3p (glyceraldehyde-3-phosphate dehydrogenase) inhibitor IAA on SICD and ROS generation in strains BY4742, *Δglr1, TRR1*_*DAmP*,_ and *Δzwf1*. Data represent the percentage of stained cells out of the total number of cells counted (100,000 cells). Cells were incubated with 2 mM of IAA for 30 min. After this, glucose was added at a final concentration of 100 mM and incubated for 1 hour at 30°C (27 °C for *TRR1*_*DAmP*_). ROS were detected by DHR fluorescence (blue). PI fluorescence was used for dead cell detection (SICD – red). Data are presented as mean±SD from 3 independent experiments.

To test this possibility, a similar IAA experiment was conducted in *Δglr1* strains. Unexpectedly, deletion of *GLR1* did not restore SICD/ROS levels (Fig. 1). That is, glutathione is not involved in ROS neutralization during SICD. This was quite surprising. The common view emphasizes the importance of glutathione in protecting the cell from oxidative stress, and it is indeed involved in numerous antioxidant reactions (Pocsi, Prade, and Penninckx 2004). However, within the cellular antioxidant machinery, it plays an auxiliary role relative to thioredoxin (Toledano *et al*. 2013).

To test the role of thioredoxin, the effect of IAA on SICD/ROS dynamics was tested in the *TRR1*_*DAmP*_ strain.Since a null TRR1 strain is non-viable, a strain with decreased TRR1 abundance via mRNA perturbation (DAmP) was used. Paradoxically, IAA still caused SICD suppression in this strain. Thus, neither glutathione nor thioredoxin is involved in SICD suppression via ROS scavenging. Two conclusions can be drawn from these data. First, our data indicate that neither the glutathione nor the thioredoxin system is responsible for ROS elimination during SICD. This suggests that ROS detected under SICD conditions are not the primary cause of cell death but rather a secondary consequence of necrotic processes. The type and localization of ROS generated during SICD are likely incompatible with classical NADPH-dependent antioxidant systems, supporting the view that SICD-associated ROS represent a marker of cellular collapse rather than a trigger of the death program. Second, it could be concluded that the non-oxidative PPP is functionally required for suppression of SICD, whereas the oxidative branch is dispensable under these conditions. Nevertheless, sufficient data confirm the possibility of independent operation of both PPP branches (Stincone *et al*. 2015).

Since SICD/ROS dynamics after 2 mM IAA treatment did not differ between BY4742 and *Δzwf1* strains, it can be assumed that glucose-6-phosphate entry into the oxidative PPP is blocked. To explain this result, the following observation must be considered: *ZWF1* expression peaks in anti-phase to the replication cluster and non-oxidative PPP genes (Clasquin *et al*. 2011), which may explain the functional silence of the oxidative branch in these processes. Consequently, GAP accumulating from GAPDH inhibition enters the PPP via the transketolase (EC 2.2.1.1)/transaldolase (EC 2.2.1.2) node. Both PPP branches produce R5P, which serves as a precursor for deoxynucleoside triphosphates (dNTPs) required for replication. SICD manifests in cells in the S-phase, when DNA synthesis is the main task. PPP activity directly affects R5P availability and thereby dNTP supply in the S-phase.Based on this, it was hypothesized that SICD/ROS suppression upon glycolysis disruption is due to carbon flux diversion into the PPP, increasing R5P production—the essential precursor for DNA synthesis. To test this, BY4742 cells were incubated under standard SICD conditions (water + 100 mM glucose) with R5P or ribose (Fig. 2). As shown in Fig. 2, ribose had no effect, whereas R5P suppressed SICD/ROS in a concentration-dependent manner, starting from 1 mM. At 5 mM, R5P inhibited SICD/ROS by more than 90%, while glucose consumption did not change compared to control. In the absence of R5P, extracellular glucose decreased by 67%, and with 5 mM R5P, by 80%, ruling out glucose uptake blockade as the cause of SICD reduction.

**Fig 2.**
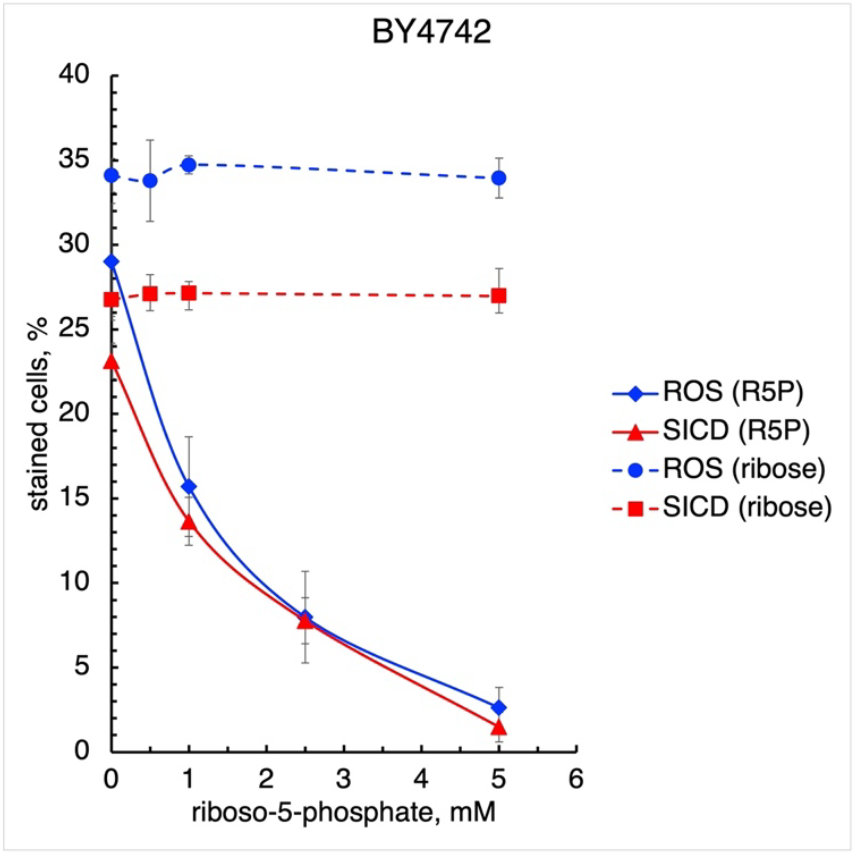
Effect of R5P and ribose on SICD and ROS generation in strain BY4742. Data represent the percentage of stained cells out of the total number of cells counted (100,000 cells). Cells were incubated with the indicated concentrations of R5P or ribose in the presence of 100 mM glucose for 1 hour at 30°C. ROS were detected by DHR fluorescence (blue). PI fluorescence was used for dead cell detection (SICD – red). Data are presented as mean±SD from 3 independent experiments.

Another possible explanation is that extracellular R5P could act as a signaling molecule, triggering antioxidant processes. To test this, the effects of other phosphorylated compounds were examined. Presence of 5 mM AMP, ADP, ATP, deoxyATP, or NADPH did not alter SICD/ROS dynamics in BY4742 (not shown), indicating that the R5P effect is specific to the phosphorylated sugar itself, not the pentose or phosphate charge (∼P^-2^). It is important to note that *S. cerevisiae* lacks specific transport systems for phosphosugars. Sugars are thought to enter unmodified and be phosphorylated intracellularly (Romano 1982; Lagunas 1993). The observed data could be explained by two mechanisms: first, R5P import via endocytosis; second, R5P dephosphorylation by constitutive acid phosphatase Pho3p (EC 3.1.3.2), ribose transport via facilitated diffusion, and re-phosphorylation by ribokinase Rbk1p (EC 2.7.1.15) (Xu *et al*. 2013; Schroeder *et al*. 2018). The *Δend3* strain, defective in endocytosis (Raths *et al*. 1993; Tuo, Nakashima, and Pringle 2013), was used to test the first mechanism. As shown in Fig. 3b, incubation with 5 mM R5P suppressed SICD/ROS to the same extent as in BY4742, ruling out endocytosis. To test the second mechanism, two approaches were used: incubation with 5 mM ribose ± Pi (to mimic Pho3p activity) in BY4742. As shown in Fig. 3a, ribose ± Pi did not affect SICD/ROS, indicating that R5P, not ribose, suppresses SICD/ROS. This effect was confirmed in the *Δpho3* strain lacking constitutive acid phosphatase (Fig. 3c).

**Fig 3.**
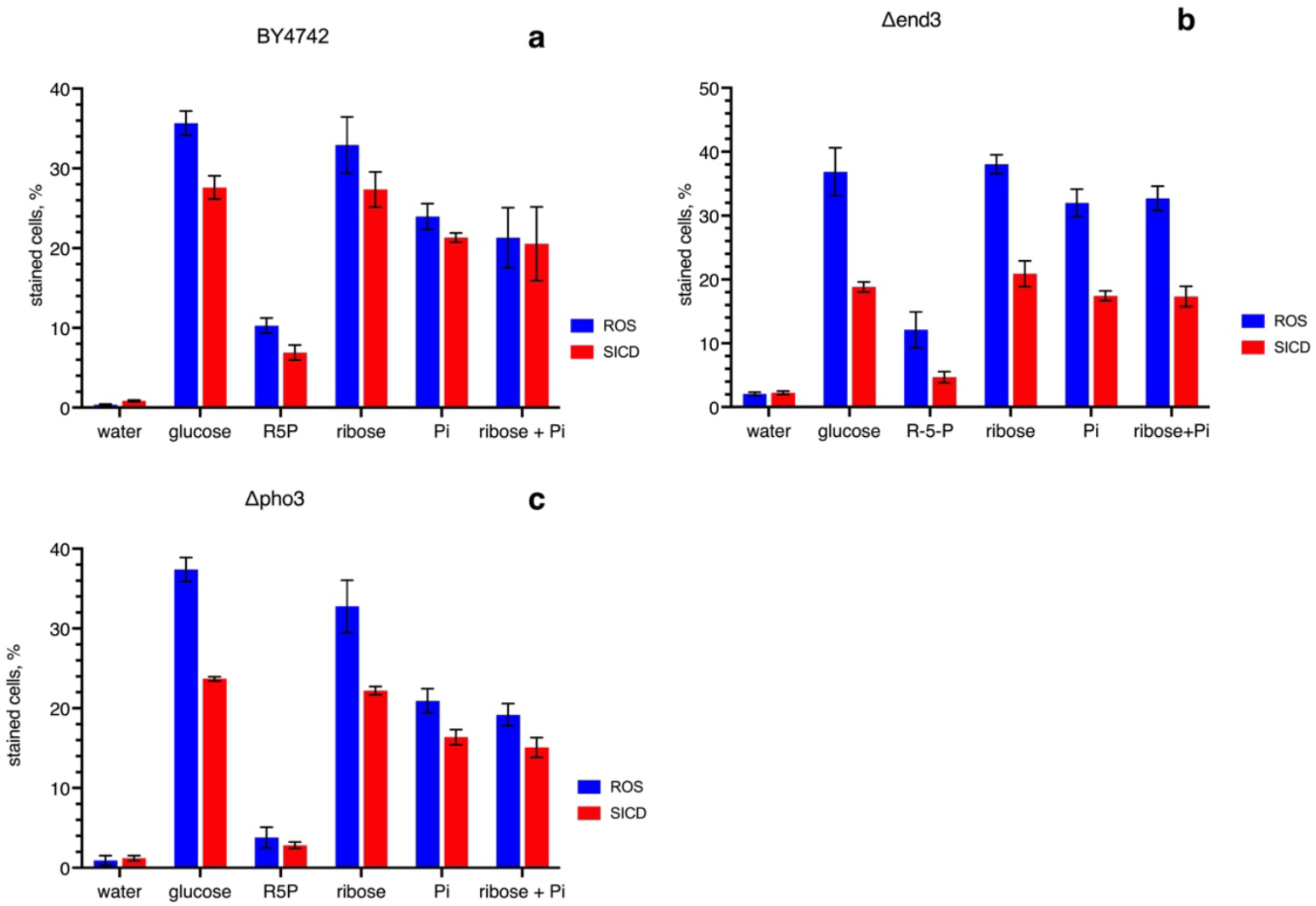
Effect of 5 mM R5P, ribose, Na_2_HPO_4,_ and ribose + Na_2_HPO_4_ on SICD and ROS generation in strains BY4742 (**a**), *Δend3* (**b**), and *Δpho3* (**c**). Data represent the percentage of stained cells out of the total number of cells counted (100,000 cells). Cells were incubated with 5 mM of R5P, ribose, Na_2_HPO_4,_ and ribose + Na_2_HPO_4_ in the presence of 100 mM glucose for 1 hour at 30°C. ROS were detected by DHR fluorescence (blue). PI fluorescence was used for dead cell detection (SICD – red). Data are presented as mean±SD from 3 independent experiments.

Thus, based on these data, R5P deficiency is the cause of SICD. Two details must be clarified. First, why ribose does not alter SICD/ROS dynamics: *S. cerevisiae* lacks specific pentose transporters. Pentoses can enter cells with low affinity via hexose transporters (Hxt and Gal2) (Young, Lee, and Alper 2010; Young *et al*. 2011). In our conditions, 100 mM glucose is actively consumed (Valiakhmetov 2025), making ribose uptake unlikely. Passive diffusion is possible but does not yield substantial R5P (Xu *et al*. 2013; Ruchala and Sibirny 2021). Second, the effect of inorganic phosphate (Pi): in stationary cells, 1 mM Pi suppressed SICD 16.7-fold and ROS 88% (Lee *et al*. 2011), but in exponentially growing cells, 5 mM Pi suppressed SICD by only 22% (BY4742), 19% (*Δend3*), and 31% (*Δpho3*), likely reflecting ionic strength effects (Avtukh *et al*. 2023).

There is currently no definitive explanation of why the PPP is not involved in glucose metabolism under SICD-inducing conditions with functional glycolysis. Intensive glycolytic flux produces fructose-2,6-bisphosphate, which activates Pfk1p (EC 2.7.1.11) (Avigad 1981). As a result, F6P preferentially enters glycolysis rather than the PPP (Yuan *et al*. 1990; Hu *et al*. 2022). Thus, under our conditions (water + 100 mM glucose), F6P entry into PPP is limited. Artificial redirection of glucose flux by glycolysis disruption (*Δhxk2, Δtps1, Δtdh3*, IAA treatment (Valiakhmetov 2025)) activates the non-oxidative PPP, producing R5P and suppressing SICD/ROS. This PPP dysfunction can be bypassed by providing exogenous R5P.

The main question regarding R5P’s effect on SICD/ROS concerns import. No specific phosphosugar transport system has been identified. Bioinformatic and screening methods also found no evidence for such systems (Andre 1995; Donzella, Sousa, and Morrissey 2023; Stanchev *et al*. 2023). The observed effect is consistent with the moonlighting function of an unknown enzyme. Moonlighting phenomena demonstrate unexpected properties of well-known enzymes and compounds (Arvizu-Rubio, García-Carnero, and Mora-Montes 2022; Curtis *et al*. 2023; Werelusz, Galiniak, and Mołoñ 2024). Why was the extracellular R5P effect not observed previously? First, the experimental system (water + 100 mM glucose) was not previously used. Second, SICD occurs only in S-phase cells, limiting detection. The combination of these factors allowed observation of R5P import resembling a salvage pathway, where the cell must choose between death and completing S-phase. Examples of salvage pathways under starvation/stress include sporulation or self-RNA utilization (Piekarska, Rytka, and Rempola 2010; Xu *et al*. 2013).

These data lead us to conclude that SICD in S-phase cells results from a deficiency of R5P required for DNA synthesis. This deficiency is a consequence of dysfunction of the PPP. One limitation of this study is that we did not directly quantify PPP intermediates. Of course, direct measurement of these metabolites would give more quantitative support. However, the present work is based mainly on functional evidence obtained by genetic perturbations and physiological readouts. The consistency between IAA-mediated glycolytic blockade, the independence from ZWF1, and the strong rescue effect of extracellular R5P allows us to infer different involvement of the two PPP branches without using metabolomic datasets.The mechanism by which R5P suppresses SICD/ROS, as well as the reasons for PPP dysfunction, require further investigation.

## References

Andre B. An overview of membrane transport proteins in Saccharomyces cerevisiae. Yeast 1995;11:1575–611.

Arvizu-Rubio VJ, García-Carnero LC, Mora-Montes HM. Moonlighting proteins in medically relevant fungi. PeerJ2022;10:e14001.

Avigad G. Stimulation of yeast phosphofructokinase activity by fructose 2,6-bisphosphate. Biochem Biophys Res Commun 1981;102:985–91.

Avtukh A, Baskunov B, Keshelava V et al. Sugar-induced cell death in the yeast S. cerevisiae is accompanied by the release of octanoic acid, which does not originate from the Fatty acid synthesis type II mitochondrial system. Applied Microbiology 2023;3:722–34.

Bertels LK, Fernandez Murillo L, Heinisch JJ. The Pentose Phosphate Pathway in Yeasts: More Than a Poor Cousin of Glycolysis. Biomolecules 2021;11, DOI: 10.3390/biom11050725.

Bidiuk VA, Alexandrov AI, Valiakhmetov AY. Extracellular pH and high concentration of potassium regulate the primary necrosis in the yeast Saccharomyces cerevisiae. Arch Microbiol 2021;204:35.

Clasquin MF, Melamud E, Singer A et al. Riboneogenesis in yeast. Cell 2011;145:969–80.

Curtis NJ, Patel KJ, Rizwan A et al. Moonlighting Proteins: Diverse Functions Found in Fungi. Journal of Fungi 2023;9:1107.

Donzella L, Sousa MJ, Morrissey JP. Evolution and functional diversification of yeast sugar transporters. Essays Biochem 2023;67:811–27.

Granot D, Dai N. Sugar induced cell death in yeast is dependent on the rate of sugar phosphorylation as determined by Arabidopsis thaliana hexokinase. Cell Death Differ 1997;4:555–9.

Granot D, Levine A, Dor-Hefetz E. Sugar-induced apoptosis in yeast cells. FEMS Yeast Res 2003;4:7–13.

Granot D, Snyder M. Glucose induces cAMP-independent growth-related changes in stationary-phase cells of Saccharomyces cerevisiae. Proc Natl Acad Sci U S A 1991;88:5724–8.

Granot D, Snyder M. Carbon source induces growth of stationary phase yeast cells, independent of carbon source metabolism. Yeast 1993;9:465–79.

Grant CM. Metabolic reconfiguration is a regulated response to oxidative stress. J Biol 2008;7:1.

Grüning N-M, Du D, Keller MA et al. Inhibition of triosephosphate isomerase by phosphoenolpyruvate in the feedback regulation of glycolysis. Open Biology 2014;4:130232.

Grüning NM, Rinnerthaler M, Bluemlein K et al. Pyruvate kinase triggers a metabolic feedback loop that controls redox metabolism in respiring cells. Cell Metab 2011;14:415–27.

Hu D, Zhang Y, Liu D et al. PFK2/FBPase-2 is a potential target for metabolic engineering in the filamentous fungus Myceliophthora thermophila. Front Microbiol 2022;13:1056694.

Kwolek-Mirek M, Maslanka R, Bednarska S et al. Disorders of Redox Homeostasis and Its Importance in Acrolein Toxicity. Int J Mol Sci 2025;26:9047.

Lagunas R. Sugar transport in Saccharomyces cerevisiae. FEMS Microbiol Rev 1993;10:229–42.

Lee YJ, Burlet E, Galiano F et al. Phosphate and succinate use different mechanisms to inhibit sugar-induced cell death in yeast: insight into the Crabtree effect. J Biol Chem 2011;286:20267–74.

Lind C, Gerdes R, Schuppe-Koistinen I et al. Studies on the mechanism of oxidative modification of human glyceraldehyde-3-phosphate dehydrogenase by glutathione: catalysis by glutaredoxin. Biochem Biophys Res Commun 1998;247:481–6.

Parbhudayal R, Cheng H-P. Sugar-induced cell death is temperature-dependent and conserved in Saccharomyces cerevisiae and Candida species. Microbiol Spectr 2025a;13:e0156825.

Parbhudayal R, Cheng H-P. Exploring Sugar-Induced Cell Death (SICD) in Yeast: Implications for Diabetes and Cancer Research. Front Cell Death 2025b;4:1470093.

Piekarska I, Rytka J, Rempola B. Regulation of sporulation in the yeast Saccharomyces cerevisiae. Acta Biochimica Polonica 2010;57:241–50.

Pocsi I, Prade RA, Penninckx MJ. Glutathione, an altruistic metabolite in fungi. Adv Microb Physiol 2004;49:1–76.

Pollak N, Dölle C, Ziegler M. The power to reduce: pyridine nucleotides--small molecules with a multitude of functions. Biochem J 2007;402:205–18.

Ralser M, Heeren G, Breitenbach M et al. Triose phosphate isomerase deficiency is caused by altered dimerization--not catalytic inactivity--of the mutant enzymes. PLoS One 2006;1:e30.

Ralser M, Wamelink MM, Kowald A et al. Dynamic rerouting of the carbohydrate flux is key to counteracting oxidative stress. Journal of Biology 2007;6:1–18.

Raths S, Rohrer J, Crausaz F et al. end3 and end4: two mutants defective in receptor-mediated and fluid-phase endocytosis in Saccharomyces cerevisiae. J Cell Biol 1993;120:55–65.

Romano AH. Facilitated diffusion of 6-deoxy-D-glucose in bakers’ yeast: evidence against phosphorylation-associated transport of glucose. J Bacteriol 1982;152:1295–7.

Ruchala J, Sibirny AA. Pentose metabolism and conversion to biofuels and high-value chemicals in yeasts. FEMS Microbiol Rev 2021;45:fuaa069.

Schroeder RY, Zhu A, Eubel H et al. The ribokinases of Arabidopsis thaliana and Saccharomyces cerevisiae are required for ribose recycling from nucleotide catabolism, which in plants is not essential to survive prolonged dark stress. New Phytol 2018;217:233–44.

Stanchev LD, Møller-Hansen I, Lojko P et al. Screening of Saccharomyces cerevisiae metabolite transporters by ^13^C isotope substrate labeling. Front Microbiol 2023;14:1286597.

Stincone A, Prigione A, Cramer T et al. The return of metabolism: biochemistry and physiology of the pentose phosphate pathway. Biological Reviews 2015;90:927–63.

Toledano MB, Delaunay-Moisan A, Outten CE et al. Functions and cellular compartmentation of the thioredoxin and glutathione pathways in yeast. Antioxid Redox Signal 2013;18:1699–711.

Tuo S, Nakashima K, Pringle JR. Role of endocytosis in localization and maintenance of the spatial markers for bud-site selection in yeast. PLoS One 2013;8:e72123.

Valiakhmetov A. Suppression of glycolysis decreases sugar-induced cell death in Saccharomyces cerevisiae. FEMS Microbiology Letters 2025;372, DOI: 10.1093/femsle/fnaf026.

Valiakhmetov AY, Kuchin AV, Suzina NE et al. Glucose causes primary necrosis in exponentially grown yeast Saccharomyces cerevisiae. FEMS Yeast Res 2019;19, DOI: 10.1093/femsyr/foz019.

Werelusz P, Galiniak S, Mołoñ M. Molecular functions of moonlighting proteins in cell metabolic processes. Biochim Biophys Acta Mol Cell Res 2024;1871:119598.

Xu Y-F, Létisse F, Absalan F et al. Nucleotide degradation and ribose salvage in yeast. Mol Syst Biol 2013;9:665.

Yoshimoto H, Ohuchi R, Ikado K et al. Sugar induces death of the bottom fermenting yeast Saccharomyces pastorianus. Journal of bioscience and bioengineering 2009;108:60–2.

Young E, Lee S-M, Alper H. Optimizing pentose utilization in yeast: the need for novel tools and approaches. Biotechnol Biofuels 2010;3:24.

Young E, Poucher A, Comer A et al. Functional survey for heterologous sugar transport proteins, using Saccharomyces cerevisiae as a host. Appl Environ Microbiol 2011;77:3311–9.

Yuan Z, Medina MA, Boiteux A et al. The role of fructose 2,6-bisphosphate in glycolytic oscillations in extracts and cells of Saccharomyces cerevisiae. Eur J Biochem 1990;192:791–5.

